# Prolonged nicotine exposure reduces aversion to the drug in mice by altering nicotinic transmission in the interpeduncular nucleus

**DOI:** 10.1101/2021.12.16.472949

**Authors:** Sarah Mondoloni, Claire Nguyen, Eléonore Vicq, Joachim Jehl, Romain Durand-de Cuttoli, Nicolas Torquet, Stefania Tolu, Stéphanie Pons, Uwe Maskos, Fabio Marti, Philippe Faure, Alexandre Mourot

## Abstract

Nicotine intake is likely to result from a balance between the rewarding and aversive properties of the drug, yet the individual differences in neural activity that control aversion to nicotine and their adaptation during the addiction process remain largely unknown. Using a two-bottle choice experiment, we observed a high heterogeneity in nicotine-drinking profiles in isogenic adult male mice, with about half of the mice persisting in consuming nicotine even at high concentrations, whereas the other half stopped consuming. We found that nicotine intake was negatively correlated with nicotine-evoked currents in the interpeduncular nucleus (IPN), and that prolonged exposure to nicotine, by weakening this response, decreased aversion to the drug, and hence boosted consumption. Lastly, using knock-out mice and local gene re-expression, we identified β4-containing nicotinic acetylcholine receptors of IPN neurons as the molecular and cellular correlates of nicotine aversion. Collectively, our results identify the IPN as a substrate of individual variabilities and adaptations in nicotine consumption.

## Introduction

Nicotine remains one of the most-widely used addictive substance in the world, and even though cigarette smoking is overall decreasing, the use of new products such as electronic cigarettes has risen dramatically in recent years ^1^. Nicotine administration induces a series of effects, ranging from pleasant (i.e. appetitive, rewarding, reinforcing, anxiolytic…) to noxious (i.e. anxiogenic, aversive…) ^2,3^. These multifaceted effects have been described both in humans and in rodents. They greatly depend on the dose of nicotine administered, show substantial inter-individual variability, and are believed to be key in the regulation of nicotine intake and in the maintenance of addiction ^2–4^. Understanding the variable effects of nicotine at the molecular and circuit levels is therefore fundamental to progress in the pathophysiology of nicotine addiction, and to develop efficient smoking-cessation therapies.

Nicotine mediates its physiological effects by activating nicotinic acetylcholine receptors (nAChRs), pentameric ligand-gated ion channels encoded by a large multigene family ^2^. There are nine nAChR α (α2-10) and three β (β2-4) subunits expressed in the brain, which can assemble to form homo-pentamers or hetero-pentamers with various localizations and functions ^5,6^. Initiation of consumption and reinforcement to nicotine involve the mesolimbic dopamine reward circuit, which originates in the ventral tegmental area (VTA) ^7^. Nicotine primarily acts on this circuit by activating α4β2 nAChRs, a receptor subtype that displays high affinity for the drug ^7–10^. Interestingly, an acute injection of nicotine also inhibits a subset of VTA dopamine neurons that project to the amygdala ^11^, and this results in elevated anxiety in mice, illustrating the heterogeneity of the brain reward circuit, and the complexity of nicotine dependence.

Another important pathway in the neurobiology of nicotine addiction is the medial habenulo-interpeduncular (MHb-IPN) axis ^12–15^. This pathway is deeply implicated in the regulation of aversive physiological states such as fear and anxiety ^16–18^. It is believed to directly mediate aversion to high doses of nicotine ^12,13^, to trigger the affective (anxiety) and somatic symptoms following nicotine withdrawal ^19–22^ and to be involved in relapse to nicotine-seeking ^23^. Strikingly, neurons of the MHb-IPN axis express the highest density and largest diversity of nAChRs in the brain, notably the rare α5, α3 and β4 subunits ^6^. These are encoded by the CHRNA5-A3-B4 gene cluster, of which some sequence variants are associated with a high risk of addiction in humans ^24,25^. The α3 and β4 subunits are virtually absent in the VTA, or in other parts of the brain. The α3β4 nAChR shows lower affinity for nicotine than the α4β2 subtype, which has contributed to the widely accepted idea that nicotine is rewarding at low doses because it activates primarily α4β2 receptors of the VTA, while it is aversive at high doses because only then does it activate α3β4 nAChRs of the MHb-IPN axis ^12,14^. Nevertheless, this hypothesis of different sensitivity to nicotine in different circuits is based on indirect evidence from c-fos quantification or brain slice physiology experiments, and does not take into account the possible adaptive changes in the circuits that result from repeated exposure to nicotine. Therefore, recording the physiological response of IPN neurons to nicotine *in vivo*, both in naive animals and after prolonged exposure to nicotine, remains a prerequisite to understand the mechanism by which smokers develop tolerance to the aversive effects of nicotine.

A distinct feature of addiction is that, overall, only some individuals lose control over their drug use, progressively shifting to compulsive drug intake ^26–30^. About a third to half of those who have tried smoking tobacco become regular users ^31^. Individual differences in the sensitivities of the VTA and MHb-IPN systems, and in their respective adaptations during chronic tobacco use, could contribute to the vulnerability to nicotine and to the severity of the addiction process ^14,32–35^. Yet, the neural mechanism that makes individuals more prone to maintain nicotine consumption versus durably stop are unclear. In addition, whether the MHb-IPN pathway respond differently to nicotine in individuals with and without a history of nicotine use is largely unknown. Here we used isogenic mice and electrophysiology (*ex vivo* and *in vivo*) to study the neuronal correlates of inter-individual variabilities in nicotine consumption behavior, and their adaptation after chronic exposure to the drug.

## Results

### Heterogeneity in nicotine consumption in isogenic wild-type mice

We used a continuous access, two-bottle choice nicotine-drinking test to assess consumption profiles in wild-type (WT) C57Bl6 male mice single-housed in their home cage (Fig. 1A). In this test, animals have continuous and concurrent access to two bottles containing a solution of either 2% saccharine (vehicle) or nicotine plus 2% saccharine (to mask the bitter taste of nicotine). After a 4-day habituation period with water in both bottles, nicotine concentration was progressively increased in one bottle across 16 days, from 10 to 200 μg/ml (4 days at each concentration), while alternating the side of the nicotine-containing solution every other day to control for side bias (Fig. 1B). Consumption from each bottle was measured every minute. We found that the daily nicotine intake increased throughout the paradigm, to stabilize at about 10 mg/kg/day on average for the highest nicotine concentration tested (Fig. 1C). Overall, the percent of nicotine consumption, i.e., the nicotine solution intake relative to the total fluid intake, was initially close to 50%, and decreased below that value for nicotine concentrations above 50 μg/ml (Fig. 1D). These results match what was observed in previous studies using C57Bl6 male mice, notably the fact that these mice rarely show over 50% nicotine consumption, whether the water is supplemented with saccharine or not ^36,37^. The decreased percent nicotine consumption observed at the population level in the course of the task suggests that mice adapt their behavior to reduce their number of visits to the nicotine-containing bottle. We indeed observed that mice rapidly (within a day) adjusted their nicotine intake when the concentration of nicotine increases (Fig. S1A), resulting in titration of the nicotine dose, as previously reported^12,34^.

**Figure 1:**
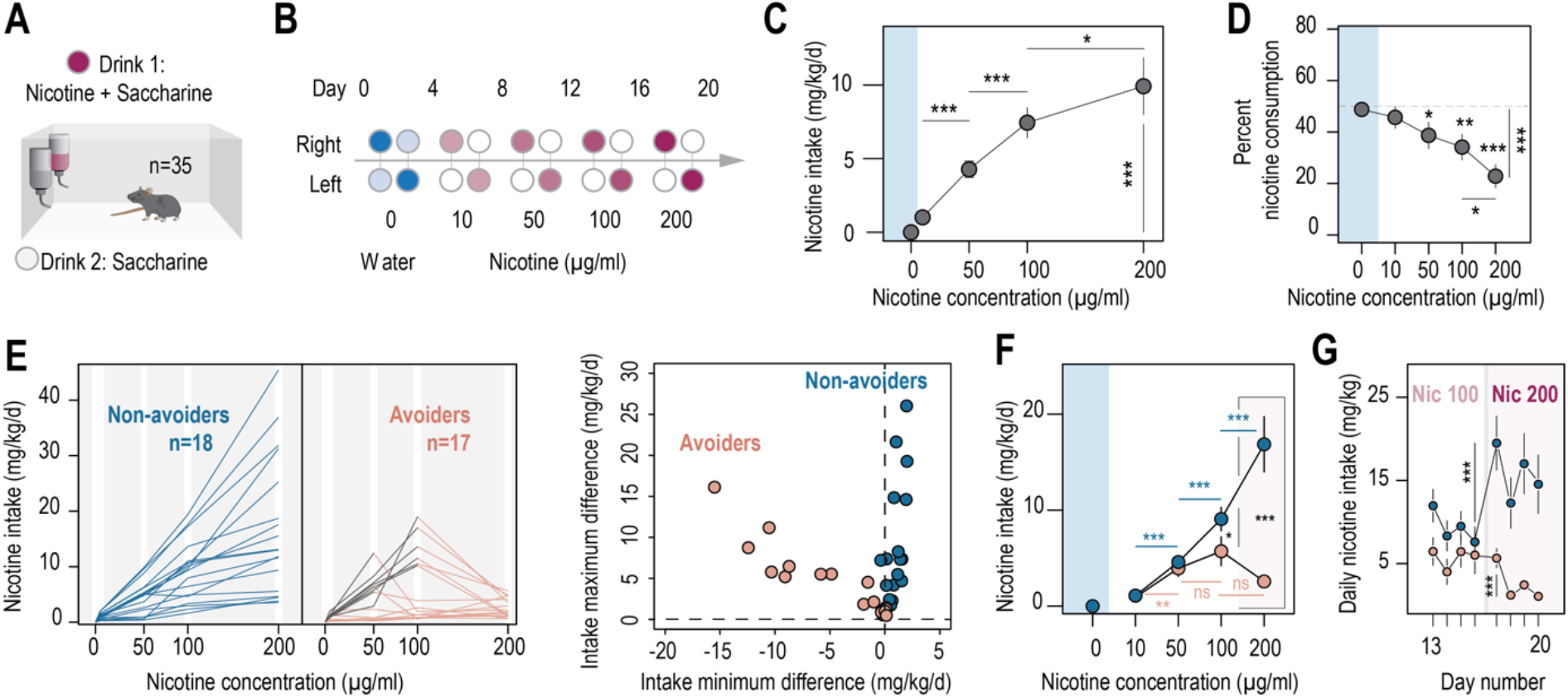
Two different profiles, avoiders and non-avoiders, emerged in WT mice subjected to a two-bottle choice nicotine-drinking test. **A.** Continuous access, two-bottle choice setup. **B.** Two-bottle choice paradigm. Each dot represents a bottle and is color-coded according to whether it contains water (blue or light blue), nicotine plus 2% saccharine (red, gradient of color intensities according to the nicotine concentration), or 2% saccharine (white) solutions. The nicotine concentration in the bottle increased progressively from 10 to 50, 100 and 200 μg/ml. Each condition lasted four days, and the bottles were swapped every other day. **C.** Nicotine intake (mg/kg/d), averaged over four days, at different nicotine concentrations (Friedman test, n = 35, df = 3, p < 0.001 and Mann-Whitney post-hoc test with Holm-Bonferroni correction). **D.** Percent nicotine consumption in WT mice for each concentration of nicotine, averaged over four days (Friedman test, n = 35, df = 4, p < 0.001 and Mann-Whitney post-hoc test with Holm-Bonferroni correction) **E.** Left, nicotine intake in individual avoiders (n = 17) and non-avoiders (n = 18). Right, minimum and maximum values of the difference in nicotine intake between two consecutive days, for each individual. **F.** Nicotine intake in avoiders and non-avoiders for each nicotine concentration, averaged over four days (Mann-Whitney comparison with a Holm-Bonferroni correction). **G.** Daily nicotine intake in avoiders and non-avoiders for nicotine 100 and 200 μg/ml (paired Mann-Whitney). Note the drop in nicotine consumption at day 17 for avoiders. In all figure panels avoiders are depicted in pinkish-orange while non-avoiders are in blue. *** p < 0.001, ** p < 0.01, * p < 0.05.

We noticed some disparity between mice and decided to analyze more precisely nicotine consumption profiles for each individual. In particular, some mice abruptly reduced their intake right after an increase in nicotine concentration in the bottle, or had very low nicotine intake throughout the entire task (intake never exceeded 2 mg/kg/day) (Fig. 1E). We classified these mice (17/35) as “avoiders”. The other half of the mice (18/35), in contrast, displayed a continuous increase in nicotine intake, or eventually reached a titration plateau in their consumption. These mice were classified as “non-avoiders”. Another way of looking at these distinct consumption profiles is to quantify the differences in intake between two consecutive days (positive differences indicate increased intake, while negative differences indicate decreased intake). The distinction between the two phenotypic groups appears clearly when we plot, for each mouse, the minimum and maximum values of the difference in intake between two consecutive days (Fig. 1E, right). The group of avoiders was characterized either by a minimum difference in intake that was negative (mice that reduce their intake abruptly), or by both a minimum and a maximum difference in intake that were close to zero (very low intake throughout the task). In contrast, the non-avoider mice were characterized by a minimum difference in intake between two consecutive days that was always positive, indicating continual increase in intake throughout the task. Overall, only a small proportion of the mice (7/35, all non-avoiders) reached a plateau in their consumption, which somewhat contrasts with the apparent titration observed at the population level, either here (Fig. 1C) or in previous studies ^12,34^. Avoiders and non-avoiders (phenotypes defined as above throughout the manuscript) showed on average similar nicotine intake for low concentrations of nicotine (10 and 50 μg/ml, p > 0.05), but while nicotine intake increased throughout the task for non-avoiders (16.9 ± 2.9 mg/kg/day for 200 μg/ml of nicotine) it dropped down to 2.6 ± 0.4 mg/kg/day for such high nicotine concentration in avoiders (Fig. 1F), and almost reached zero over the last three days (Fig. 1G). The percent nicotine consumption was fairly constant throughout the task in non-avoiders, whereas in avoiders, it drastically decreased as nicotine concentration increased, to approximate zero at the end of the task (Fig. S1B). We then compared the level of aversion produced by nicotine in avoiders and non-avoiders, with that produced by quinine, a notoriously bitter molecule. We found that all of the naive mice actively avoided the quinine-containing solution, and showed a percent quinine consumption close to zero, as observed with nicotine for avoiders, but not for non-avoiders (Fig. S1C). Together, these results suggest that avoiders display strong aversion to nicotine.

### The concentration of nicotine triggering aversion differs between mice

Owing to the oral nature of the test, it may be difficult for mice to associate consumption in a particular bottle with the physiological effects (whether positive or negative) of nicotine. We thus systematically analyzed the percent nicotine consumption throughout the task in individual mice, to examine their variability in choice patterns and behavioral alterations. We found that some non-avoiders actively tracked the side associated with nicotine when bottles were swapped (e.g. mouse #1 in Fig. 2A), indicating strong preference for the bottle containing nicotine and active consumption. Other non-avoiders displayed a strong side preference and never alternated drinking side (e.g. mouse #2 in Fig. 2A), and hence consumed nicotine in a more passive fashion. In contrast, all avoider mice (n = 17) displayed avoidance of the nicotine-containing solution, whether they initially tracked the nicotine solution (e.g. mice #3 and 4 in Fig. 2A) or not (e.g. mice #5 and 6 in Fig. 2A). To quantify the evolution of nicotine consumption at the individual level throughout the task, and to better take into account the passive consumption behavior of some of the mice, we mapped each profile in a pseudo-ternary plot where the base represents the nicotine consumption index (from 0 to 100%), while the top apex represents 100% side bias (Fig. 2B). Such a ternary representation enables us to graphically distinguish between mice that actively track the nicotine bottle (bottom right apex, 100% nicotine consumption index), mice that actively avoid nicotine (bottom left apex, 0% nicotine consumption index), and mice that have a strong side bias (top apex), and to calculate the shortest distance for each mouse to each of the three apices. In addition, by representing the trajectory for each individual from the water condition to the 200 μg/ml nicotine condition, this plot can be used to reveal and quantify behavioral adaptations (or lack thereof) in each individual. Overall, we found that the behavior of non-avoiders was on average fairly consistent throughout the task, i.e. their distance from the three apices was not really impacted by the modifications in nicotine concentration (Fig. 2C and Fig. S2), whether they consumed nicotine actively or passively. In contrast, the behavior of avoiders was highly nicotine concentration-dependent, with mice going further away from the 100% nicotine apex as nicotine concentration increases (Fig. 2C and Fig. S2A-B). Aversion to nicotine in avoider mice mainly occurred at the transition from 100 to 200 μg/ml, but some mice displayed aversion at concentrations as low as 10 μg/ml (Fig. 1E, 2A and 2C). Together, these results show that most mice can actively track or avoid nicotine, indicating that they can discriminate nicotine from the control solution. Avoiders started actively avoiding the nicotine-containing bottle once a specific drug concentration was reached, suggesting the existence of a threshold at which nicotine aversion is triggered, which apparently differs between mice.

**Figure 2:**
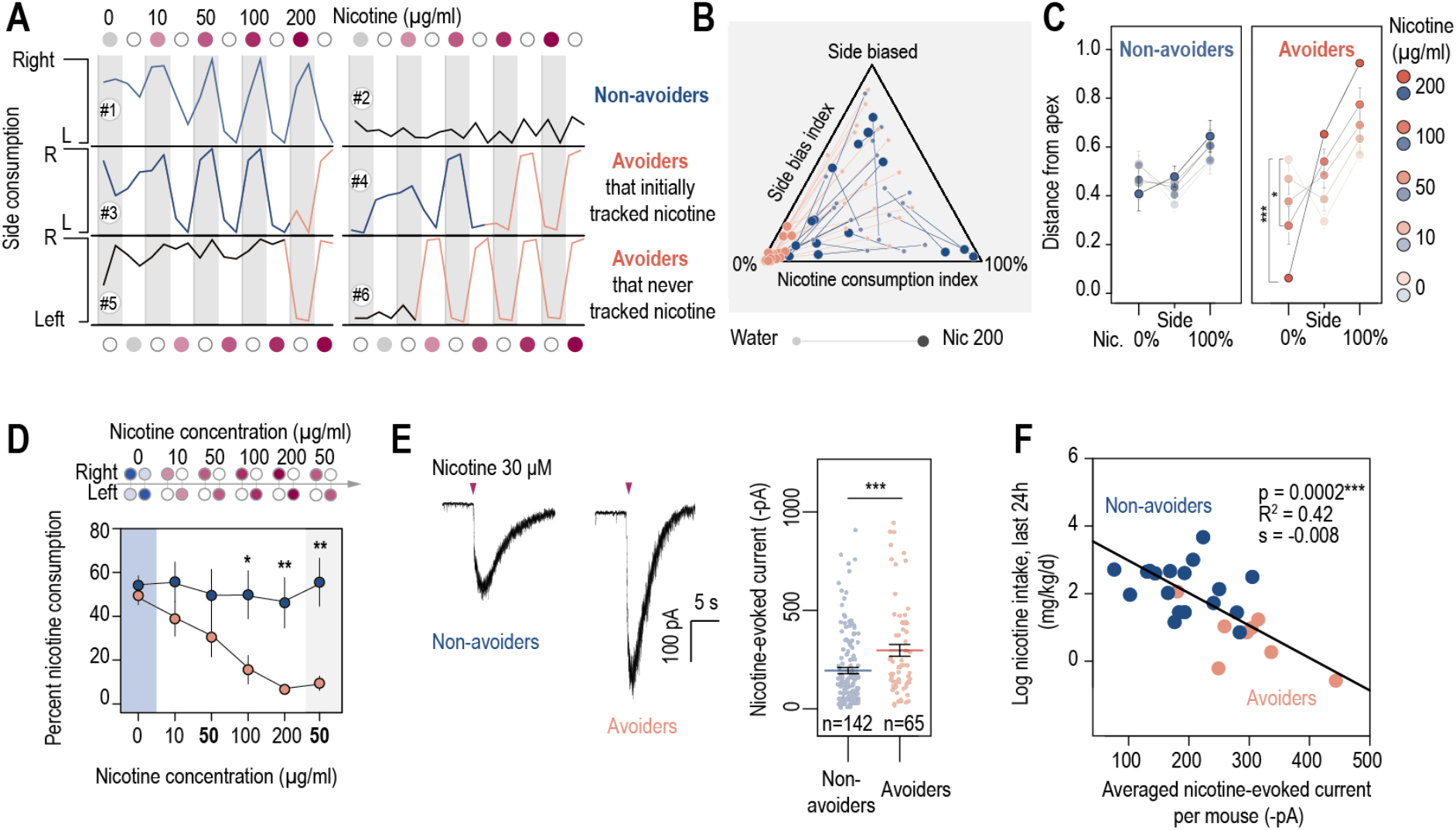
Nicotine intake was negatively correlated with the amplitude of the response to nicotine in IPN neurons. **A.** Representative examples of choice behaviors (% consumption on the right vs. left bottle) in WT mice when the right-hand side bottle contains either nicotine + saccharine (red dots, grey stripe) or saccharine only (white dots, white stripes). Mice are same as in Fig. 1. **B.** Pseudo-ternary diagram representing, for each individual (18 non-avoiders and 17 avoiders, see Fig. 1), its nicotine consumption index over its side bias index. Bottom left apex: 0% nicotine consumption (0%); Bottom right apex: 100% nicotine consumption (100%); Top apex: 100% side preference (Side biased, i.e. mice that never switch side). Small dots correspond to the habituation period (water vs. water) while bigger dots correspond to the condition with 200 μg/ml of nicotine in one bottle. Note how all avoider mice end up in the bottom left apex (0% nicotine consumption) at the end of the task. **C.** Average distance from the three apices for each condition in the task (0, 10, 50, 100 and 200 μg/ml of nicotine, color-coded from light to dark), for avoiders and non-avoiders. Only avoider mice significantly changed their drinking strategy as nicotine concentration increased (paired Mann-Whitney test with Holm-Bonferroni correction). **D.** Average percent nicotine consumption for avoider and non-avoider mice for each concentration of nicotine. The 50 μg/ml nicotine solution was presented a second time to the mice, at the end of the session, for four days. (Mann-Whitney test, Holm-Bonferroni correction, p(50 Nic) = 0.04, p(100 Nic) = 0.005, p(200 Nic) = 0.004). **E.** Representative (left) and average (right) currents recorded in voltage-clamp mode (−60 mV) from IPN neurons of non-avoiders (blue, n = 142 neurons from 18 mice, I = −194 ± 15 pA) and avoiders (red, n = 65 neurons from 8 mice, I = −297 ± 30 pA) following a puff application of nicotine (30 μM, 200 ms). Avoiders presented greater nicotine-evoked currents than non-avoiders (Mann-Whitney test, p = 1.4e-11). **G.** Correlation between the dose consumed (log scale, over the last 24 hours prior to the recording) and the averaged nicotine evoked-current per mouse (-pA). In all figure panels avoiders are depicted in pinkish-orange and non-avoiders in blue. *** p<0.001, ** p<0.01, * p<0.05.

Do avoiders learn to keep away from the nicotine-containing solution, or do they just rapidly react to the nicotine concentration in the bottle to adjust their daily intake? To answer this question, we added at the end of the two-bottle choice task, i.e. after the 200 μg/ml nicotine concentration, a condition with a low concentration of nicotine (50 μg/ml) for four days. We chose 50 μg/ml of nicotine because avoiders and non-avoiders initially displayed comparable nicotine intake (Fig. 1F) and percent consumption (Fig. S1B) at this concentration. We hypothesized that if avoiders increased their percent nicotine consumption at the 200 to 50 μg/ml transition, it would indicate a rapid adjustment to the concentration proposed, so to maintain their level of intake constant. In contrast, if avoiders maintained a steady, low percent nicotine consumption, it would indicate that nicotine aversion persists, independently of the dose. We indeed found that lowering nicotine concentration from 200 to 50 μg/ml did not increase percent nicotine consumption in avoiders (Fig. 2D), at least for the four days mice were subjected to this concentration, suggesting aversion learning for nicotine, and a behavioral adaptation that leads to near complete cessation of nicotine consumption.

### Nicotine consumption negatively correlated with the amplitude of nicotine-evoked currents in the IPN

We then investigated the neural correlates of such aversion to nicotine. We hypothesized that the IPN, which is involved in nicotine aversion and in negative affective states ^3,38,39^, might be differently activated by nicotine in avoiders and non-avoiders. Therefore, we used whole-cell patch-clamp recordings in brain slices to assess, at completion of the two-bottle choice task, the functional expression level of nAChRs in IPN neurons. We recorded neurons from the dorsal and rostral IPN because these neurons have high nAChR density ^14,15,40,41^. To record nicotine-evoked currents, we used a local puff application of nicotine at a concentration (30 μM) close to the EC50 for heteromeric nAChRs ^42^. We found that the amplitude of nicotine-evoked currents was higher in IPN neurons of avoider mice than in non-avoiders (Fig. 2E). Moreover, we found a negative correlation between the average amplitude of nicotine-evoked current in IPN neurons, and nicotine consumption (measured over the last 24 hours prior to the patch-clamp recording, Fig. 2F), suggesting that nicotine consumption in mice is negatively linked to the amplitude of the response to nicotine in IPN neurons.

### Chronic nicotine treatment alters both nicotinic signaling in the IPN and nicotine consumption

It is still unclear at this stage whether chronic nicotine exposure progressively alters the response of IPN neurons to the drug (non-avoiders being further exposed to high nicotine doses than avoiders) or whether an intrinsic difference pre-exists in avoiders and non-avoiders. To determine the effect of chronic nicotine exposure on nAChR current levels in IPN neurons, we passively and continuously exposed mice to nicotine for 4 weeks, using subcutaneously implanted osmotic minipumps. The concentration of nicotine in the minipump (10 mg/kg/day) was chosen to match the average voluntary nicotine intake in the two-bottle choice task (Fig. 1C). We then recorded from acute brain slices and found that indeed, prolonged exposure to nicotine reduced the amplitude of nicotine-evoked currents in the IPN of these mice compared to control mice treated with saline (Fig. 3A). These results are consistent with the reduced current amplitudes observed in mice that underwent the two-bottle choice task, compared to naive mice in their home cage (Fig S3A). Because IPN neurons are mostly silent in brain slices, and in order to preserve the entire circuitry intact, we decided to perform juxtacellular recordings of IPN neurons *in vivo*, and to characterize their response (expressed in % of variation from baseline) to an intravenous (i.v.) injection of nicotine (30 μg/kg). To our knowledge, no description of *in vivo* recordings of IPN neurons, and thus no criteria for identification, have been reported as yet, hence we solely considered neurons that were labelled *in vivo* with neurobiotin and confirmed to be within the IPN for the analysis. We found that nicotine i.v. injections in naive WT mice induced an increase in the firing rate of IPN neurons compared to an injection of saline, and that this acute effect of nicotine was reduced after the prolonged (4 weeks) passive exposure of mice to the drug (Fig.3B). Some of the IPN neurons, however, responded to nicotine by decreasing their firing rate, and the degree of inhibition was lower in the group exposed to chronic nicotine (Fig. S3B). Together, these *ex vivo* and *in vivo* recordings demonstrate that prolonged exposure to nicotine markedly reduces nicotine-evoked responses in mouse IPN neurons.

**Figure 3:**
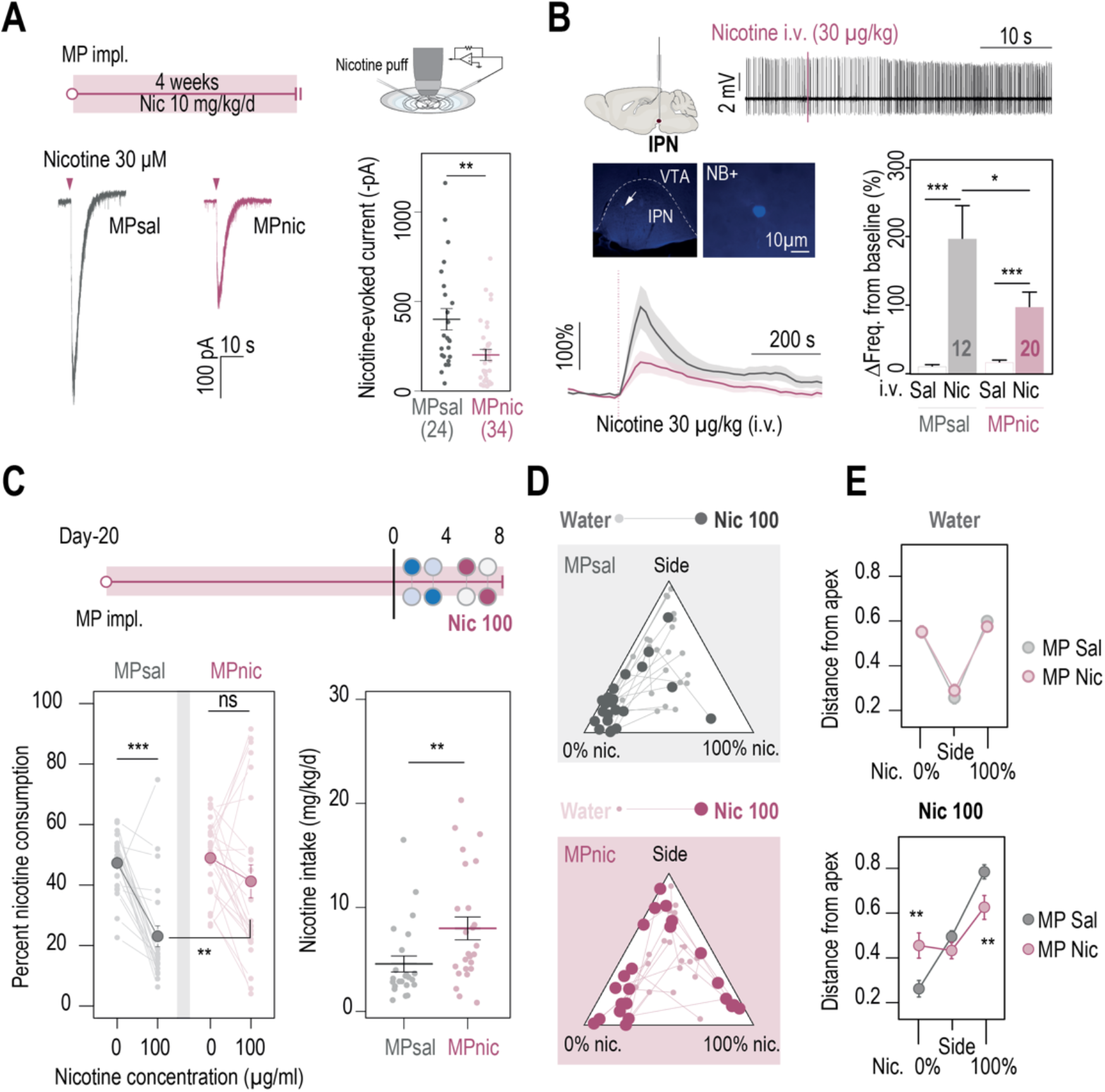
Chronic nicotine treatment altered both nAChR expression levels in the IPN and nicotine intake in WT mice. **A.** Top, passive nicotine treatment protocol. Mice were implanted subcutaneously with an osmotic minipump (MP) that continuously delivers 10 mg/kg/d of nicotine. After 4 weeks of treatment, nicotine-evoked responses in IPN neurons were recorded in whole-cell voltage-clamp mode (−60 mV) from IPN slices. Bottom, representative recordings (left) and average current amplitudes (right) following a puff application of nicotine (30 μM, 200 ms) in IPN neurons of mice treated with either saline (n = 24 neurons from 3 mice, I = −401 ± 59 pA) or nicotine (n = 34 neurons from 4 mice, I = −202 ± 37 pA). Nicotine treatment reduced the amplitude of nicotine-evoked currents in IPN neurons (Mann-Whitney test, p = 0.001). **B.** *In vivo* juxtacellular recordings of nicotine-evoked responses in IPN neurons of saline# and nicotine-treated animals. Top, representative electrophysiological recording of an IPN neuron, during an i.v. injection of nicotine (30 μg/kg). Middle, post-recording identification of neurobiotin-labeled IPN neurons by immunofluorescence. Bottom, average time course and average amplitude of the change in firing frequency from baseline after an i.v. injection of saline and nicotine (30 μg/kg), for IPN neurons from saline# and nicotine-treated mice. Right, firing rate variation from baseline induced by nicotine or saline injection in IPN neurons from saline-(n = 6) or nicotine-treated (n = 13) animals. Responses were decreased by chronic exposure to nicotine (p = 0.035, Mann-Whitney test). All neurons were confirmed to be located within the IPN using juxtacellular labeling with neurobiotin. **C.** Top, modified two-bottle choice protocol used to evaluate the impact of a long-term exposure to nicotine on drug intake. Mice were implanted subcutaneously with a minipump that delivered 10 mg/kg/d of nicotine continuously, for 20 days before performing the modified two-bottle choice task. After four days of water vs. water habituation, mice were directly exposed to a high concentration of nicotine (100 μg/ml). Bottom, percent nicotine consumption and nicotine intake at 0 and 100 μg/ml of nicotine, for mice under a chronic treatment of nicotine or saline. The saline-treated group displayed a decrease in percent nicotine consumption (n = 23, from 47.3 ± 2.0% to 23.0 ± 3.4%, p = 1.7e-05, Mann-Whitney paired test), but not the the nicotine-treated group (n = 25, from 48.9 ± 2.4 to 41.2 ± 5.5%, p = 0.16, Mann-Whitney paired test). Overall, the saline-treated group displayed a lower percent nicotine consumption (p = 0.003, Mann Whitney) and lower nicotine intake than the nicotine-treated group (p = 0.004, Mann-Whitney). **D.** Pseudo-ternary diagrams representing each saline# and nicotine-treated mouse for its nicotine consumption index over its side bias index. Small dots correspond to the habituation period (water vs. water) and bigger dots to the condition with 100 μg/ml of nicotine in one bottle. **E.** Average distance from each apex in the water vs. water (top) and water vs. nicotine 100 μg/ml conditions (bottom). Saline-treated, but not nicotine-treated mice developed a strategy to avoid nicotine (p_Sacc_ = 0.013, p_Side_ = 0.27 p_Nic_ = 0.013, Mann-Whitney test with Holm-Bonferroni correction). In all figure panels nicotine-treated animals are displayed in red and saline-treated (control) animals in grey.

To verify the hypothesis that a modification in the cholinergic activity of IPN neurons impacts nicotine aversion, we did a series of experiments. Firstly, we evaluated the consequence of prolonged nicotine exposure on nicotine consumption. Mice were implanted with an osmotic minipump to passively deliver nicotine and, after 20 days, were subjected to a modified two-bottle task that consisted in a rapid presentation to a high concentration (100 μg/ml) of nicotine (Fig. 3C). We chose this protocol to avoid the confounding effects of a gradual exposure to nicotine, and to evoke strong aversion in mice. We found that indeed, mice pretreated with saline (controls) showed low percent nicotine consumption (Fig. 3C), which may indicate aversion to such high nicotine concentration. In contrast, mice pretreated with nicotine did not decrease on average their percent consumption when nicotine was introduced in the bottle, suggesting that they may have developed tolerance for the aversive effects of nicotine. When looking at the data day by day, we observed that control mice abruptly decreased their percent consumption when nicotine was introduced, while for nicotine-treated animals the decrease was more gradual over the four days (Fig. S3C). Overall, this resulted in greater nicotine intake for the group treated with nicotine than for the group treated with saline (Fig. 3C). When focusing on individuals, we observed that a single saline-pretreated mouse (1/23) increased its percent consumption when nicotine was introduced in the task, while the great majority of the mice actively avoided nicotine (close to the 0% Nicotine apex). In contrast, a marked proportion of the mice treated with nicotine (8/25) increased their percent consumption when nicotine was introduced (Fig. 3D, p-value = 0.037 Chi-squared). The two groups were identical in the water/water session (Fig. 3E, top). However, in the water/nicotine session, mice pretreated with nicotine showed a greater distance from the 0% nicotine consumption apex, and a shorter distance from the 100% nicotine consumption apex than mice pretreated with saline (Fig. 3E, bottom), indicating decreased aversion and increased preference for nicotine in the nicotine-pretreated group. Altogether, these electrophysiological and behavioral data demonstrate that prolonged exposure to nicotine both decreases nicotine efficacy in the IPN, and also decreases aversion to nicotine in individuals, resulting in increased consumption of the drug. Yet, chronic nicotine can produce adaptations in other brain circuits, and whether the neurophysiological changes observed in the IPN are causally related to the variations in aversion sensitivity remains to be demonstrated.

### Nicotine avoidance involves β4*nAChRs

We turned to mutant mice deleted for the gene encoding the nAChR β4 subunit (β4^−/−^ mice), because of the strong and restricted expression of this subunit in the MHb-IPN pathway ^32,43,44^. We found that β4^−/−^ mice displayed both greater percent nicotine consumption and greater nicotine intake than WT animals, with minimal concentration-dependent change in percent nicotine consumption (Fig. 4A). When looking at individuals, we observed both active and passive nicotine-drinking profiles in β4^−/−^ mice, as already observed in WT mice. Strikingly however, none of the β4^−/−^ mice (0/13) showed aversion-like behavior at high nicotine concentration, which contrasts with the high proportion of avoiders in WT animals (17/35, Fig. 4B, Fig. S4A, p =0.04 Pearson’s Chi squared with Yates’continuity correction). WT mice showed a strong nicotine concentration-dependent adaptation in their behavior, while β4^−/−^ mice had a more consistent behavior throughout the task (Fig. 4C and Fig. S4B, C).

**Figure 4:**
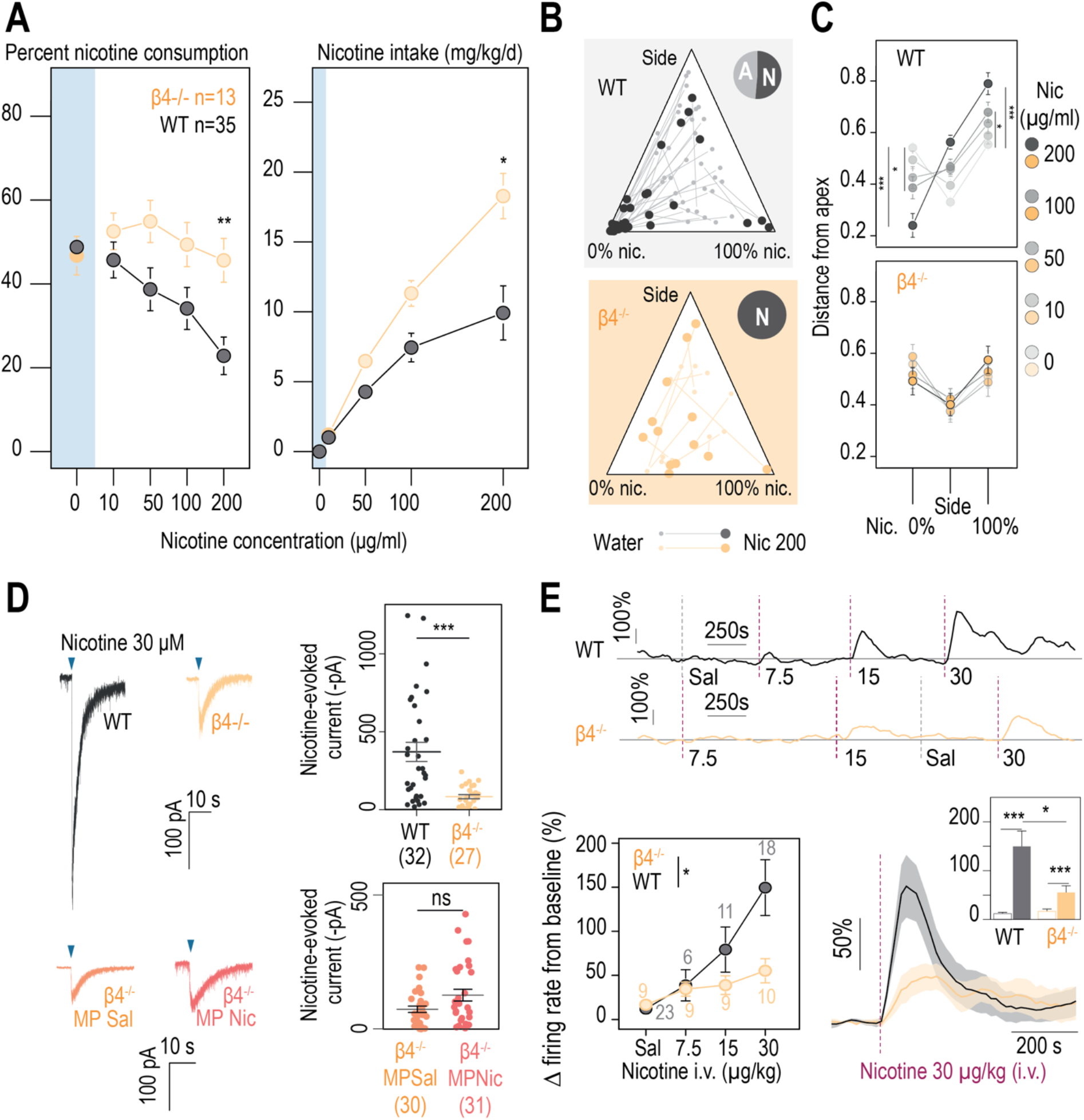
β4-containing nicotinic receptors are essential for triggering nicotine aversion in mice. **A.** Left, average percent nicotine consumption in WT and β4^−/−^ mice for each concentration of nicotine in the two-bottle choice task. WT mice had lower percent nicotine consumption than β4^−/−^ mice (Mann-Whitney test with Holm-Bonferroni correction). WT mice decreased their percent nicotine consumption throughout the task (Friedman test, n = 35, df = 4, p < 0.001 and Mann-Whitney post-hoc test with Holm-Bonferroni correction) while β4^−/−^ mice displayed a stable percent nicotine consumption (Friedman test, n = 13, df = 4, p = 0.11). Right, average nicotine intake (mg/kg/d) in β4^−/−^ and WT mice for the different concentrations of nicotine (Friedman test, n = 35, df = 3, p < 0.001 and Mann-Whitney post-hoc test with Holm-Bonferroni correction). β4^−/−^ mice consumed more nicotine than WT mice (Mann-Whitney test). **B.** Ternary diagram representing each WT and β4^−/−^ individual for its nicotine consumption index over its side bias index. Small dots correspond to the habituation period (water vs. water) and bigger dots to the condition with 200 μg/ml of nicotine in one bottle. Inserts: pie charts illustrating the proportion of avoiders (A, light grey) and non-avoiders (N, dark grey) for each genotype at the end of the task. Note the absence of avoiders in β4^−/−^ mice. **C.** Average distance from each apex at 0, 10, 50, 100 and 200 μg/ml of nicotine (paired Mann-Whitney test with Holm-Bonferroni correction, p(Sacc 0-200) = 0.0002, p(Sacc 0-100) = 0.03; p(Nic 0-200) = 0.0002, p(Sacc 0-100) = 0.03). **D.** Left, representative currents following a puff application of nicotine (30 μM, 200 ms) in IPN neurons from naive WT and β4^−/−^ mice, or from saline-treated (orange) and nicotine-treated (dark orange) β4^−/−^ mice. Right, average nicotine-evoked currents recorded in IPN neurons from naïve WT (n = 32 neurons from 5 mice, I = −370 ± 61 pA) and β4^−/−^ (n = 27 neurons from 4 mice, I = −83 ± 13 pA) mice, and from β4^−/−^ chronically treated with either saline (Sal, n = 30 neurons from 6 mice, I = −72 ± 11 pA) or nicotine (Nic, n = 31 neurons from 5 mice, I = −123 ± 21 pA). β4^−/−^ mice presented a large decrease in nicotine-evoked currents (Mann-Whitney test, p = 4e-05). Nicotine treatment did not alter nicotine-evoked currents in IPN neurons of β4^−/−^ mice (Mann-Whitney test, p = 0.15). **E.** Juxtacellular recordings of nicotine-evoked responses in IPN neurons in naive WT and β4^−/−^ mice. Top, representative example of the variation in firing frequency of an IPN neuron, following repeated i.v. injections of nicotine at 7.5, 15 and 30 μg/kg, in WT and β4^−/−^ mice. Bottom left, dose-dependent change in firing rate from baseline following i.v. injections of nicotine (p < 0.05). All recorded neurons were neurobiotin-labelled to confirm their location within the IPN. Bottom right, average nicotine-evoked responses at 30 μg/kg of nicotine in IPN neurons from WT and β4^−/−^ mice. Insert, average amplitude of the change in firing frequency from baseline after an i.v. injection of saline and nicotine (30μg/kg), for IPN neurons from WT (n = 18) and β4^−/−^ (n = 7) mice. Nicotine-induced responses were smaller in β4^−/−^ than in WT mice (p < 0.001, Mann Whitney test). In all figure panels WT animals are depicted in grey and β4^−/−^ mice in yellow. *** p < 0.001, ** p < 0.01, * p < 0.05.

To verify that responses to nicotine were affected in IPN neurons of β4^−/−^ mice, we performed whole-cell patch-clamp recordings. We found that the amplitude of nicotine-evoked currents in the IPN was on average three-fold lower in β4^−/−^ than in WT mice (Fig. 4D), confirming that β4*nAChRs are the major receptor subtype in the IPN. Furthermore, prolonged nicotine treatment had no significant effect on the amplitude of nAChR currents in these knock-out mice (Fig. 4D), suggesting that the downregulation observed after chronic nicotine treatment in the IPN of WT mice mainly affects β4*nAChRs. We then used *in vivo* juxtacellular recordings, and performed dose-response experiments (7.5 - 30 μg/kg) in order to assess the role of β4*nAChRs in the response to different doses of nicotine. In WT mice, nicotine i.v. injections resulted in a dose-dependent increase in IPN neuron activity (Fig. 4E). In β4^−/−^ mice, responses to nicotine were of smaller amplitude, especially for the highest dose of nicotine tested (Fig. 4E), further demonstrating the important role of β4*nAChRs in the response of the IPN to nicotine. In both WT and β4^−/−^ mice, we observed a population of neurons that decreased their firing rate in a dose-dependent manner, yet with no difference in the amplitude of the response between the two genotypes (Fig. S4D-F), suggesting that β4*nAChR are mainly involved in the increase, but not in the decrease, of neuronal activity in response to nicotine injection. Collectively, our results in β4^−/−^ mice demonstrate the key role of the nAChR β4 subunit in signaling aversion to nicotine, and its predominant function in the activation of the IPN by nicotine.

### β4*nAChRs of the IPN are critically involved in nicotine aversion

β4 nAChRs are enriched in the IPN, yet they are also expressed to some extent in other brain regions. Hence, to directly implicate β4*nAChRs of IPN neurons in nicotine aversion, and more generally in nicotine consumption, we targeted re-expression of β4 in the IPN specifically, using lentiviral vectors in β4^−/−^ mice (KO-β4^IPN^ mice, Fig. 5A). Mice transduced with eGFP (KO-GFP^IPN^ mice) were used as controls. Proper transduction in the IPN was verified using immunohistochemistry after completion of the two-bottle choice task (Fig. 5A), and mice with expression of GFP in the VTA were excluded from analyses. Transduction of β4, but not of GFP alone, in the IPN increased the amplitude of nicotine-evoked currents (Fig. 5B) and restored levels found in WT animals (U test, p = 0.6, Fig. S5A). We compared nicotine intake in the two groups of mice in the two-bottle choice task. We found that re-expression of β4 in the IPN of β4^−/−^ mice decreased nicotine intake compared to the group of mice transduced with eGFP in the IPN (Fig. 5C). Overall, nicotine intake was similar in WT and in KO-β4^IPN^ animals (p>0.5 for all concentrations), demonstrating the causal role of β4 nAChRs of IPN neurons in nicotine consumption behaviors. At the individual level, the proportion mice that avoided nicotine at 200 μg/kg was very low for KO-GFP^IPN^ control mice (2/18), but greater for KO-β4^IPN^ mice (8/17, Fig. 5D, p = 0.04 Chi squared). The behavior of KO-GFP^IPN^ was steady throughout the task, whereas it was highly nicotine concentration-dependent for KO-β4^IPN^ mice (Fig. 5E and Fig. S5B,C), as already observed with WT mice. Mice with strong percent nicotine consumption were only found in the KO-GFP^IPN^ control group. Collectively, these results show that selective re-expression of β4 in the IPN of β4^−/−^ mice rescued aversion for nicotine, and highlight the specific role of β4*nAChRs of IPN neurons in signaling aversion to nicotine, and in the control of nicotine intake.

**Figure 5:**
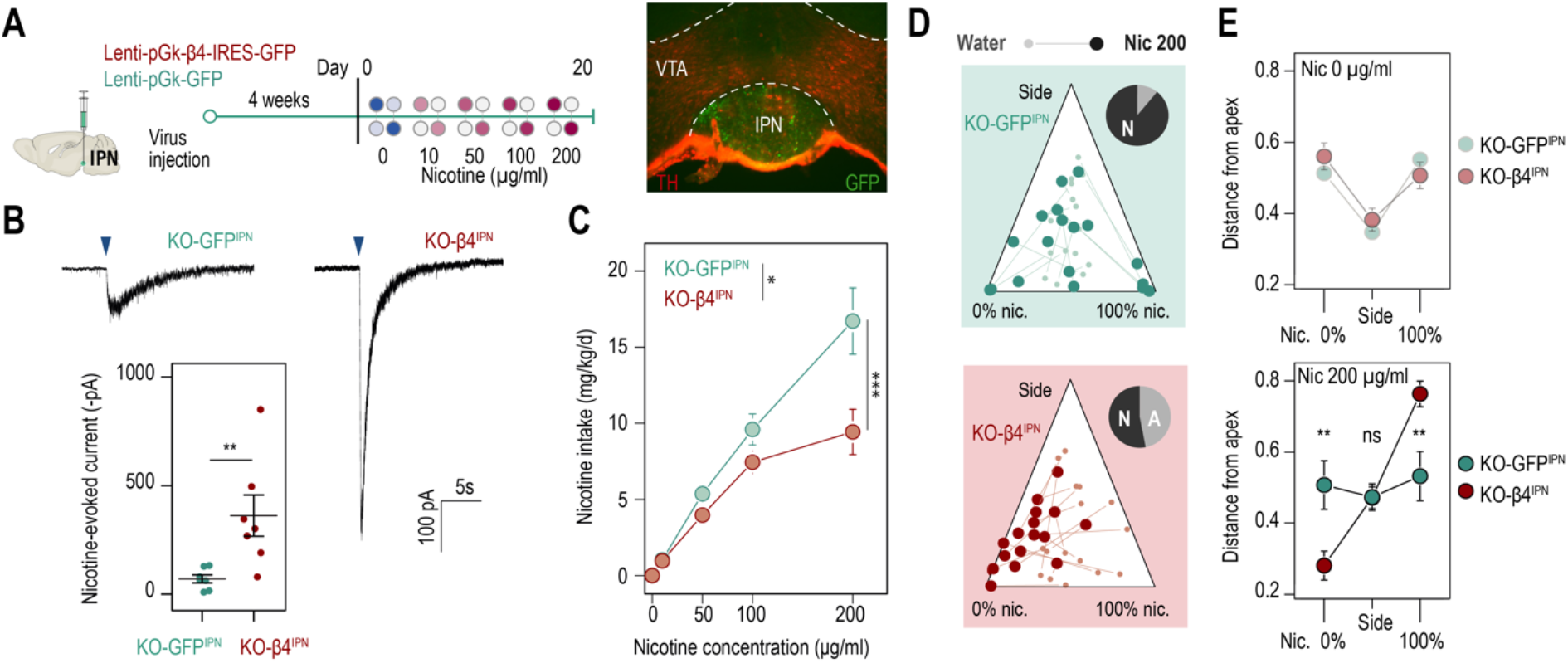
β4-containing nAChRs of the IPN are involved in the control of nicotine consumption and in aversion to nicotine in mice. **A.** Protocol: stereotaxic transduction of the β4 subunit together with GFP (or GFP alone in control mice) in the IPN of β4^−/−^ mice, and subsequent two-bottle choice task. Right: coronal section highlighting proper viral transduction of lenti-pGK-β4-IRES-GFP in the IPN. **B.** Validation of the re-expression using whole-cell patch-clamp recordings. Representative currents and average responses following a puff application of nicotine (30 μM, 200 ms) on IPN neurons from β4^−/−^ mice transduced in the IPN with either lenti-pGK-β4-IRES-GFP (KO-β4^IPN^, n = 7 neurons from 2 mice, I = −362 ± 95 pA) or lenti-pGK-GFP (KO-GFP^IPN^, n = 7 neurons from 1 mouse, I = −71 ± 18 pA; Mann-Whitney test, p = 0.004). **C.** Average nicotine intake was lower in KO-β4^IPN^ than in KO-GFP^IPN^ (two-way repeated measure; ANOVA: genotype x dose interaction, F[3, 99] = 6.3, ***p < 0.001; main effect of dose, F[3, 99] = 69.1 ***p < 0.001, effect of genotype, F[1, 33] = 6.637, *p = 0.015). **D.** Ternary diagram representing each β4^−/−^ mouse, transduced with either β4 or GFP, and illustrating its nicotine consumption index over its side bias index. Small dots correspond to the habituation period (water vs. water) and bigger dots to the condition with 200 μg/ml of nicotine in one bottle. Inserts: pie charts illustrating the proportion of avoiders (A, light grey) and non-avoiders (N, dark grey) for each condition at the end of the task. **E.** Average distance from each apex during the two-bottle choice task at 0 and 200 μg/ml of nicotine, for KO-β4^IPN^ and KO-GFP^IPN^ mice (p_Sacc_ = 0.007, p_Nic_ = 0.008, Mann-Whitney test with Holm-Bonferroni correction). In all figure panels KO-β4^IPN^ mice are depicted in red and KO-GFP^IPN^ mice (controls) in green. *** p < 0.001, ** p < 0.01, * p < 0.05.

## Discussion

We used a two-bottle choice paradigm to assess inter-individual differences in nicotine consumption in mice, and to evaluate how pre-exposure to nicotine modifies drug taking. Oral self-administration is a classical method for chronic nicotine administration as it provides rodents with *ad libitum* access to nicotine, likely mimicking administration in human smokers, while minimizing stress from handling ^45^. We observed that, on average, WT mice titrate their intake to achieve a consistent nicotine dose, in agreement with previous reports ^12,34,35^, and discovered that mutant mice lacking the nAChR β4 subunit did not, resulting in greater nicotine intake, notably at high nicotine concentrations in the drinking solution. These behavioral results are in agreement with the greater intracranial self-administration observed at high nicotine doses in these mice ^33^ and, conversely, with the results obtained with transgenic TABAC mice overexpressing the β4 subunit at endogenous sites, which avoid nicotine and consequently consume very little ^13,33^. Nevertheless, it should be noted that conflicting results have also been reported regarding the role of β4 nAChRs in nicotine consumption. Notably, intravenous self-administration of nicotine is lower in β4^−/−^ mice despite a higher sensitivity of the VTA to nicotine in these mice ^32^, and self-administration is higher in TABAC mice despite reduced nicotine-induced activation of the VTA ^46^. The increased consumption at high nicotine concentration reported here for β4^−/−^ mice resembles what was observed in mutant mice either lacking the α5 subunit ^12^ or with low levels of the α3 subunit ^47^, likely because the α3, α5, and β4 nAChR subunits, which are encoded by the same gene cluster, co-assemble in brain structures, notably the MHb-IPN pathway, to produce functional heteromeric nAChRs that contribute to the control of nicotine intake.

In the two-bottle choice nicotine-drinking test, a percent nicotine consumption below 50% is usually interpreted as a sign of aversion to the drug ^13,37^. However, it may rather indicate a process of regulation of nicotine consumption, through which mice adapt to an increase in nicotine concentration by reducing the volume of nicotine intake at each visit. At low nicotine concentrations, percent nicotine consumption never really exceeded 50% in average, as already seen in previous studies ^37^, yet very few mice readily avoided the nicotine-containing bottle, and many mice did track nicotine, which would rather indicate an actual appetite (as opposed to an aversion) for the drug. As nicotine concentration increases, the percent nicotine consumption dropped to about 20% on average at the end of the task (Fig. 1D), yet it was associated with an increase in nicotine intake (in mg/kg/day, Fig. 1C). Hence the nicotine preference score alone may not be sufficient to determine whether nicotine is aversive or not. We believe that, instead, aversion to the drug may be better defined by a constantly low consumption, or a sudden drop in nicotine intake, as seen here with the group of avoider mice (Fig. 1E-G).

One important limitation of population-level analyses is that they greatly limit the ability to examine inter-individual differences in drug taking behaviors. It is indeed increasingly acknowledged that in mice, as in humans, there is a substantial variability in the susceptibility for developing drug use disorders ^27–29,48–51^. Yet, it is still unclear why some individuals are more susceptible than others to become regular users. Here we inspected drinking profiles in individual isogenic mice, and discovered large inter-individual differences in nicotine vulnerability: about half of the WT mice, the avoiders, durably quit nicotine at a certain concentration, whereas the other half, the non-avoiders, continued consumption even at high concentration of nicotine, classically described as aversive ^3^. Avoiders displayed variable concentration thresholds required for triggering aversion, and some of them even developed aversion at the beginning of the two-bottle choice task, when concentrations of nicotine were still low. This finding, which challenges the popular idea that aversion is only triggered by high doses of nicotine ^3,12–14^, is somewhat supported by our electrophysiogical data. Indeed, by directly measuring the response of IPN neurons to different doses of nicotine *in vivo*, we show that concentrations of nicotine as low as 7.5 μg/kg can engage the IPN circuitry, suggesting that nicotine-induced signaling can emerge in this structure even at low drug concentration. Non-avoiders (WT and β4^−/−^ mice) may retain an aversion threshold, but which would exceed the highest concentration tested here. Importantly, very few mice showed what could be considered as titration (plateaued consumption), emphasizing the needs to consider individual behavior, as opposed to group behavior, in addiction research.

We also discovered that the functional expression level of β4-containing nAChRs in the IPN underlies these different sensitivities to the aversive properties of nicotine. Indeed, inter-individual variability for nicotine aversion was nearly eliminated in β4^−/−^ mice, none of which quit drinking nicotine, and was restored after selective re-expression of the β4 subunit in the IPN. This selective re-expression experiment discards the possibility that different sensitivities to the bitter taste of nicotine solutions explain the different drinking profiles of avoiders and non-avoiders. Strikingly, we observed a negative correlation between nicotine consumption and the response of the IPN to the drug: mice consuming large amounts of nicotine displayed lower nicotine-evoked currents in the IPN than mice consuming small amounts of nicotine, which displayed large currents. The expression level of β4-containing nAChRs in the IPN may thus determine the level of aversion to the drug, and consequently its intake. We suggest that β4-containing nAChRs, by engaging the IPN circuitry, initiate a primary response to nicotine that, if above a certain threshold, will trigger acute aversion to the drug, thus impacting the balance between drug reward and aversion to limit drug consumption. In line with this, it was found that pharmacological or optogenetic stimulation of the MHb-IPN pathway could directly produce aversion^14,15,34^, while pharmacological inactivation of this pathway increases nicotine intake ^12^. From our results, we further suggest that the aversion produced by nicotine is not just an acute response to the dose that has just been ingested. Aversion can also be lasting for days, as evidenced by the fact that mice persist in avoiding the nicotine-containing bottle even after the concentration has been reduced to a dose they used to ingest. This echoes observations made in humans, in which the unpleasant initial responses to cigarettes is associated with a reduced likelihood of continued smoking ^52^. Such a sustained aversive reaction to nicotine was conditioned, in mice, by nicotine itself, and required β4-containing nAChRs of the IPN for its onset, but most likely involves other brain circuits for its persistence in the long-term. Identifying the molecular and cellular mechanism of long-term aversion to nicotine in mice will be instrumental to progress in our understanding of human dependence to tobacco.

We did not observe major differences in nicotine intake between avoiders and non-avoiders at the beginning of the two-bottle choice experiment, for low concentrations of nicotine (< 100 μg/ml), but cannot completely rule out pre-existing inter-individual differences that may explain the opposite trajectories taken by the two groups. Indeed, the mice used in this study were isogenic, yet epigenetic changes during development or differences in social status, which are known to affect brain circuits and individual traits ^53^, may affect the responses of the IPN to nicotine. It is indeed tempting to speculate that external factors (e.g. stress, social interactions…) that would affect the expression level of β4-containing nAChRs in the IPN, will have a strong impact on nicotine consumption. In addition to pre-existing differences, history of nicotine use could produce long-lasting molecular and cellular adaptations in the IPN circuitry that may alter nicotine aversion and consumption. In the VTA, chronic nicotine upregulates the number of β2-containing receptors at the cell surface ^54,55^. We discovered that prolonged nicotine exposure had the opposite effect on β4-containing nAChRs of the IPN: it downregulated and/or desensitized these receptors, as evidenced by the decreased response to nicotine both *ex vivo* and *in vivo*. This effect seems to be specific to β4-containing nAChRs, since chronic nicotine had minimal consequence on the residual IPN nAChR current in β4^−/−^ mice. These results may appear at odds with the recent report of increased nAChR currents in IPN slices of nicotine-treated mice ^56^. However, differences in the effective agonist concentrations used in the two studies (30 μM of nicotine with local puff here, *vs.* 50 μM of caged-nicotine, but effective concentration probably much lower after photo-uncaging in ^56^) may explain this discrepancy. Indeed, by using a lower effective agonist concentration, Arvin et al. ^56^ were possibly not recruiting the low-affinity β4-containing receptors, which are the receptor subtypes we found to be downregulated by chronic nicotine exposure (Fig. 4D). Whatever the reason behind these opposite results is, our behavioral data in nicotine-treated WT and β4^−/−^ mice completely match: both displayed reduced responses to nicotine (*ex vivo* and i*n vivo)* in IPN neurons, as well as increased nicotine intake compared to naive, WT animals. We suggest that long-term exposure to nicotine decreases the likelihood to reach the threshold at which mice develop aversion to the drug, which ultimately leads to increased drug consumption. In other words, nicotine intake history weakens the ability of nicotine to induce aversion in mice. In most nicotine replacement therapies, such as gums or patches, nicotine is slowly administered over prolonged periods of time, to supposedly attenuate the negative emotional reactions elicited by nicotine withdrawal ^57^. However, our data indicate that mice under prolonged nicotine administration will also develop tolerance to the aversive effects of nicotine, highlighting the necessity to develop alternative medical approaches.

## Materials and Methods

### Animals

Eight to sixteen-week old wild-type C57BL/6J (Janvier labs, France) and Acnb4 knock-out (β4^−/−^) mice (Pasteur Institute, Paris) ^58^ were used for this study. β4^−/−^ mice were backcrossed onto C57BL/6J background for more than twenty generations. Mice were maintained on a 12h light-dark cycle. All experiments were performed in accordance with the recommendations for animal experiments issued by the European Commission directives 219/1990, 220/1990 and 2010/63, and approved by Sorbonne Université.

### Two-bottle choice experiment

Mice single-housed in a home cage were presented with two bottles of water (Volvic) for a habituation period of 4 days. After habituation, mice were presented with one bottle of saccharine solution (2%, Sigma Aldrich) and one bottle of nicotine (free base, Sigma Aldrich) plus saccharine (2%) solution diluted in water (adjusted to pH ~7.2 with NaOH). Unless otherwise noted, four different concentrations of nicotine were tested consecutively (10, 50, 100 and 200 μg/ml) with changes in concentration occurring every 4 days. For the two-bottle aversion task, a single nicotine concentration (100 μg/ml) was used after the habituation period. Bottles were swapped every other day to control for side preference. The drinking volume was measured every minute with an automated acquisition system (TSE system, Germany). Nicotine intake was calculated in mg of nicotine per kilogram of mouse body weight per day (mg/kg/d). To minimize stress from handling, mice were weighed every other day, since we found their weigh to be sufficiently stable over two days. Percent nicotine consumption was calculated as the volume of nicotine solution consumed as a percentage of the total fluid consumed. Mice showing a strong side bias (preference <20% or >80%) in the habituation period were not taken into account for the analyses.

For the pseudo-ternary plot analyses, we determined the percent nicotine consumption on the left-hand side (%c1) and the percent nicotine consumption on the right-hand side (%c2), for each nicotine concentration and each animal. We then calculated the nicotine consumption and side bias indexes, by plotting the minimum min(%c1, %c2) against the maximum max(%c1, %c2), and used a 90° rotation to obtain the pseudo-ternary plot. In this plot, the three apices represent mice that avoid nicotine on both sides (0% nic.), mice that track nicotine on both sides (100% nic.), and finally mice that drink solely one side (side biased).

### Prolonged treatment with nicotine

Osmotic minipumps (2004, Alzet minipump) were implanted subcutaneously in 8-week-old mice anesthetized with isoflurane (1%). Minipumps continuously delivered nicotine (10 mg/kg/d) or saline (control) solution with a rate of 0.25 μl/h during 4 weeks.

### Brain slice preparation

Mice were weighed and then anaesthetized with an intraperitoneal injection of a mixture of ketamine (150 mg/kg, Imalgene 1000, Merial, Lyon, France) and xylazine (60 mg/kg, Rompun 2%, Bayer France, Lyon, France). Blood was then fluidized by an injection of an anticoagulant (0.1mL, heparin 1000 U/mL, Sigma) into the left ventricle, and an intra-cardiac perfusion of ice-cold (0-4°C), oxygenated (95% O_2_/5% CO_2_) sucrose-based artificial cerebrospinal fluid (SB-aCSF) was performed. The SB-aCSF solution contained (in mM): 125 NaCl, 2.5 KCl, 1.25 NaH_2_PO_4_, 5.9 MgCl_2_, 26 NaHCO_3_, 25 sucrose, 2.5 glucose, 1 kynurenate (pH 7.2). After rapid brain sampling, slices (250 μm thick) were cut in SB-aCSF at 0-4°C using a Compresstome slicer (VF-200, Precisionary Instruments Inc.). Slices were then transferred to the same solution at 35°C for 10 min, then moved and stored in an oxygenated aCSF solution at room temperature. The aCSF solution contained in mM: 125 NaCl, 2.5 KCl, 1.25 NaH_2_PO_4_, 2 CaCl_2_, 1 MgCl_2_, 26 NaHCO_3_, 15 sucrose, 10 glucose (pH 7.2). After minimum 1h of rest, slices were placed individually in a recording chamber at room temperature and infused continuously with aCSF recording solution at a constant flow rate of about 2 ml/min.

### Ex vivo patch-clamp recordings of IPN neurons

Patch pipettes (5-8 MΩ) were stretched from borosilicate glass capillaries (G150TF-3, Warner instruments) using a pipette puller (Sutter Instruments, P-87, Novato, CA) and filled with a few microliters of an intracellular solution adjusted to pH 7.2, containing (in mM): 116 K-gluconate, 20 HEPES, 0.5 EGTA, 6 KCl, 2 NaCl, 4 ATP, 0.3 GTP and 2 mg/mL biocytin. Biocytin was used to label the recorded neurons. The slice of interest was placed in the recording chamber and viewed using a white light source and a upright microscope coupled to a Dodt contrast lens (Scientifica, Uckfield, UK). Neurons were recorded from the dorsal (IPDL) and rostral (IPR) parts on the IPN. Whole-cell configuration recordings of IPN neurons were performed using an amplifier (Axoclamp 200B, Molecular Devices, Sunnyvale, CA) connected to a digitizer (Digidata 1550 LowNoise acquisition system, Molecular Devices, Sunnyvale, CA). Signal acquisition was performed at 10 kHz, filtered with a lowpass (Bessel, 2 kHz) and collected by the acquisition software pClamp 10.5 (Molecular Devices, Sunnyvale, CA). Nicotine tartrate (30 μM in aCSF) was locally and briefly applied (200 ms puffs) using a puff pipette (glass pipette ~3 μm diameter at the tip) positioned about 20-30 μm from the soma of the neuron. The pipette was connected to a Picospritzer (PV-800 PicoPump, World Precision Instruments) controlled with pClamp to generate transient pressure in the pipette (~2 psi). Nicotine-evoked currents were recorded in voltage-clamp mode at a membrane potential of −60 mV. All electrophysiology traces were extracted and pre-processed using Clampfit (Molecular Devices, Sunnyvale, CA) and analyzed with R.

### In vivo electrophysiology

Mice were deeply anesthetized with chloral hydrate (8%, 400 mg/kg) and anesthesia was maintained throughout the experiment with supplements. Catheters were positioned in the saphenous veins of the mice to perform saline or nicotine intravenous injections. Nicotine hydrogen tartrate salt (Sigma-Aldrich) was dissolved in 0.9% NaCl solution and pH was adjusted to 7.4. The nicotine solution was injected at a dose of 7.5, 15 and 30 μg/kg. Borosilicate glass capillaries (1.5 mm O.D. / 1.17 mm I.D., Harvard Apparatus) were pulled using a vertical puller (Narishige). Glass pipettes were broken under a microscope to obtain a ~1 μm diameter at the tip. Electrodes were filled with a 0.5% NaCl solution containing 1.5% of neurobiotin tracer (AbCys) yielding impedances of 6-9 MΩ. Electrical signals were amplified by a high-impedance amplifier (Axon Instruments) and supervised through an audio monitor (A.M. Systems Inc.). The signal was digitized, sampled at 25 kHz and recorded on a computer using Spike2 (Cambridge Electronic Design) for later analysis. IPN neurons were recorded in an area corresponding to the following stereotaxic coordinates (4-5° angle): 3.3 - 3.6 mm posterior to bregma, 0.2 - 0.45 mm from medial to lateral and 4.3 - 5 mm below the brain surface. A 5 min-baseline was recorded prior to saline or nicotine i.v. injection. For the dose-response experiments, successive randomized injections of nicotine (or saline) were performed, interspaced with sufficient amount of time (> 10 min) to allow the neuron to return to its baseline.

### Stereotaxic viral injections

8-week old mice were injected in the IPN with a lentivirus that co-expresses the WT β4 subunit together with eGFP (or only eGFP for control experiments) under the control of the PGK promoter. Lentiviruses were produced as previously described ^7^. For viral transduction, mice were anaesthetized with a gas mixture containing 1-3% isoflurane (IsoVet®, Pyramal Healthcare Ltd., Nothumberland, UK) and placed in a stereotactic apparatus (David Kopf Instruments, Tujunga, CA). Unilateral injections (0.1μl/min) of 1μl of a viral solution (Lenti.pGK.β4.IRES.eGFP, titer 150 ng/μl of p24 protein; or Lenti.pGK.eGFP, titer 75 ng/μl of p24 protein) were performed using a cannula (diameter 36G, Phymep, Paris, France). The cannula was connected to a 10 μL Hamilton syringe (Model 1701, Hamilton Robotics, Bonaduz, Switzerland) placed in a syringe pump (QSI, Stoelting Co, Chicago, IL, USA). Injections were performed in the IPN at the following coordinates (5° angle): from bregma ML - 0.4 mm, AP - 3.5 mm, and DV: - 4.7 mm (according to Paxinos & Franklin). Electrophysiological recordings were made at least 4 weeks after viral injection, the time required for the expression of the transgene, and proper expression was subsequently checked using immunohistochemistry.

### Immunocytochemical identification

Immunostaining was performed as described in ^10^, *with the following* primary antibodies: anti-tyrosine hydroxylase 1:500 (anti-TH, Sigma, T1299) and chicken anti-eYFP 1:500 (Life technologies Molecular Probes, A-6455). Briefly, serial 60 μm-thick sections of the midbrain were cut with a vibratome. Slices were permeabilized for one hour in a solution of phosphate-buffered saline (PBS) containing 3% bovine serum albumin (BSA, Sigma; A4503). Sections were incubated with primary antibodies in a solution of 1.5 % BSA and 0.2 % Triton X-100 overnight at 4°C, washed with PBS and then incubated with the secondary antibodies for 1 hour. The secondary antibodies were Cy3-conjugated anti-mouse (1:500 dilution) and alexa488-conjugated anti-chicken (1:1000 dilution) (Jackson ImmunoResearch, 715-165-150 and 711-225-152, respectively). For the juxtacellular immunostaining, the recorded neurons were identified with the addition of AMCA-conjugated streptavidin (1:200 dilution) in the solution (Jackson ImmunoResearch). Slices were mounted using Prolong Gold Antifade Reagent (Invitrogen, P36930). Microscopy was carried out either with a confocal microscope (Leica) or with an epifluorescence microscope (Leica), and images were captured using a camera and analyzed with ImageJ.

### Statistical analysis

All statistical analyses were computed using R (The R Project, version 4.0.0). Results were plotted as a mean ± s.e.m. The total number (n) of observations in each group and the statistics used are indicated in figure legends. Classical comparisons between means were performed using parametric tests (Student’s T-test, or ANOVA for comparing more than two groups when parameters followed a normal distribution (Shapiro test P > 0.05)), and non-parametric tests (here, Mann-Whitney or Friedman) when the distribution was skewed. Multiple comparisons were corrected using a sequentially rejective multiple test procedure (Holm-Bonferroni correction). All statistical tests were two-sided. P > 0.05 was considered not to be statistically significant.

## Acknowledgments

The authors would like to thank Ines Centeno-Lemaire (Sorbonne Université, Paris, France) for her help with behavioral tests, Jean-Pierre Hardelin (ESPCI, Paris, France) for critical reading of the manuscript, and the animal facility at Institut de Biology Paris Seine (IBPS, Paris, France).

## Author contribution

Conceptualization: SM, PF and AM.

Methodology: SM, PF and AM.

Software: NT and SM.

Validation: SM, PF and AM.

Formal analysis: SM.

Investigation: SM, CN, EV, JJ, RDC, ST and FM.

Resources: SP and UM.

Writing - original draft: AM.

Writing - review and editing: SM, PF and AM.

Visualization: SM.

Supervision: PF and AM.

Funding acquisition: PF and AM.

## Data and material availability

All data needed to evaluate the conclusions in the paper are present in the paper and/or the Supplementary materials.

