## Supplementary material for "Prolonged nicotine exposure reduces aversion to the drug in mice by altering nicotinic transmission in the interpeduncular nucleus"

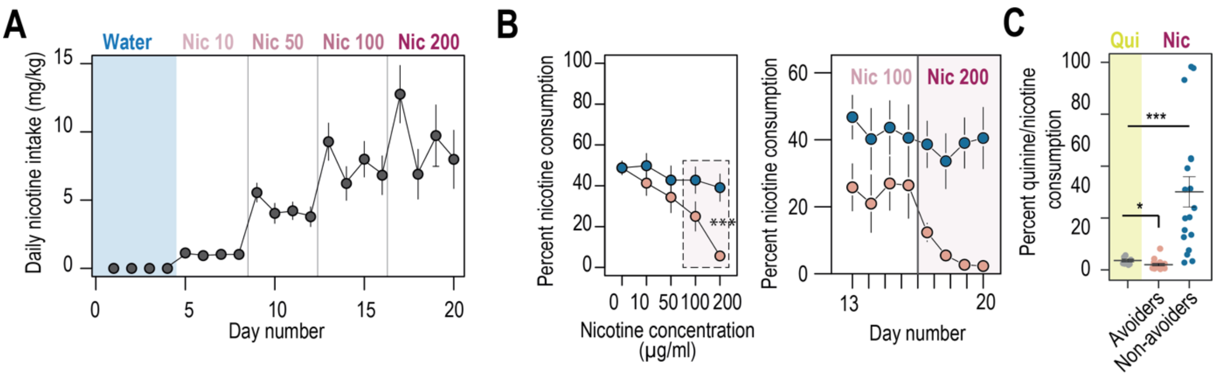


**FIGURE SUPP 1**

**Figure S1: Nicotine intake and consumption profiles in avoiders and non-avoiders. A.** Nicotine intake measured every day (mg/kg/d) for WT mice in the two-bottle choice task (n = 35). **B.** Percent nicotine consumption for avoider and non-avoider mice for each concentration of nicotine, averaged over four days for the entire session (left) or depicted each day for the last 8 days (right). **C.** Percent quinine consumption in WT animals (n = 8 mice, 4.3 +/- 0.6 %), compared to percent nicotine consumption in avoiders (n = 17 mice, 2.3 +/- 0.59 %, Mann-Whitney p-value= 0.033) and non-avoiders (n = 18 mice, 41.6 +/- 8.9 %, Mann-Whitney p-value= 0.0003). *** $p < 0.001$, ** $p < 0.01$, * $p < 0.05$.

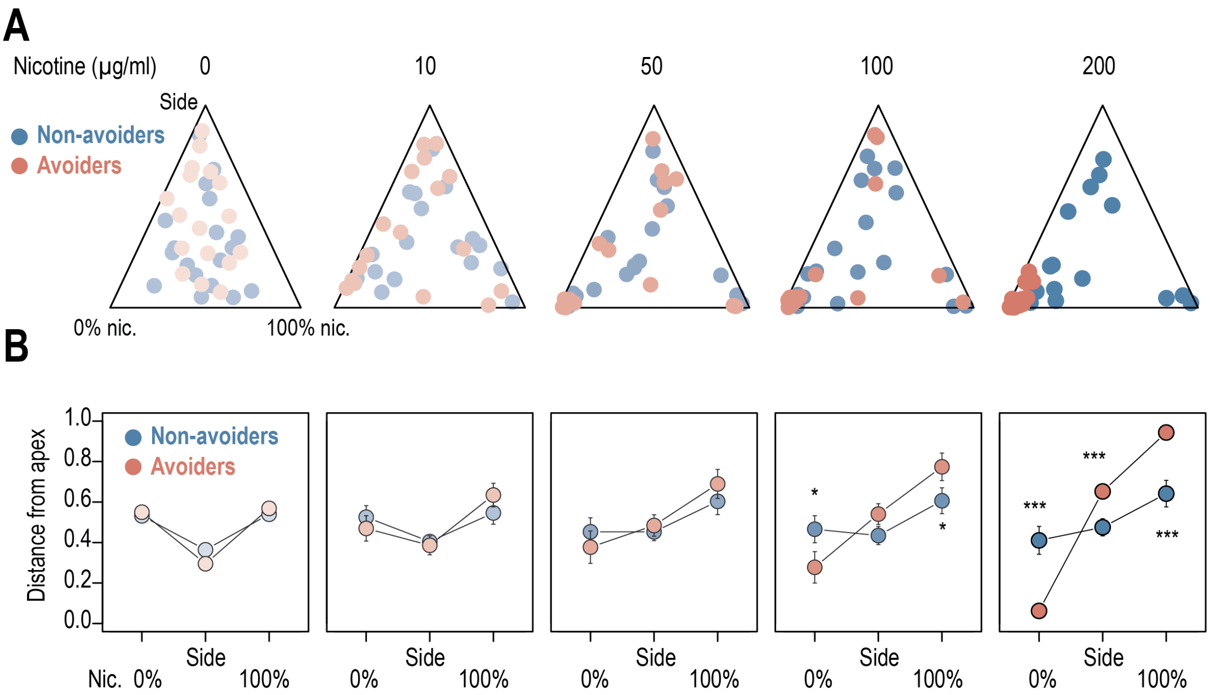


### FIGURE SUPP 2

**Figure S2: individual adaptations for avoiders and non-avoiders. A.** Ternary representations for avoiders (red) and non-avoiders (blue), illustrating their nicotine consumption index over their side bias index, as nicotine concentration increases during the task. Note how all avoiders end up, at the end of the task, in the bottom left apex corresponding to 0% nicotine consumption. **B.** Distance from each of the three apices for avoiders (red) and non-avoiders (blue) as nicotine concentration increases during the task. Note how the behavior of avoiders is highly nicotine-concentration dependent (Mann-Whitney, Nic 100 µg/ml : $p(Sacc) = 0.018$; $p(Nic) = 0.011$; Nic 200 µg/ml : $p(Sacc) < 0.001$, $p(Side) < 0.001$, $p(Nic) < 0.001$). *** $p < 0.001$, ** $p < 0.01$, * $p < 0.05$.

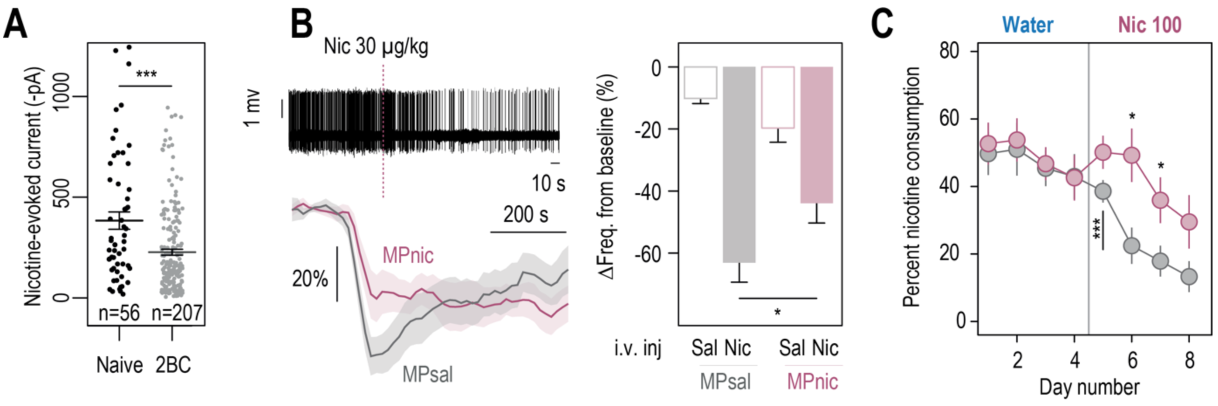


### FIGURE SUPP 3

**Figure S3: Physiological and behavioral adaptations following chronic nicotine treatment. A.** Nicotine-induced current (puff application, 30 µM, 200 ms) recorded in voltage-clamp mode (-60 mV) from IPN neurons of naive mice kept in their home-cage (black, n = 56 neurons from 8 mice, I = -384 ± 43 pA) and of mice that underwent the two-bottle choice task (2BC, grey, n = 207 neurons from 26 mice, I = -227 ± 15 pA). Nicotine-induced currents were of smaller amplitude in 2BC mice than in naive mice (Mann-Whitney, p = 0.0002). **B.** *In vivo* juxtacellular recordings of nicotine-evoked responses in nicotine-inhibited IPN neurons of saline- and nicotine-treated animals. Top left, representative electrophysiological recording of an IPN neuron, during an i.v. injection of nicotine (30 µg/kg). Bottom left, average time course and amplitude of the change in firing frequency from baseline after an i.v. injection of saline and nicotine (30 µg/kg), for IPN neurons of saline- (n = 12neurons from 7 mice) and nicotine-treated animals (n = 13neurons from 10mice). Right, responses were on average lower after chronic exposure to nicotine (p < 0.05). **C.** Daily percent nicotine consumption for saline- and nicotine-treated mice, for each day in the two-bottle choice task. In saline-treated mice, avoidance to nicotine appears from the second nicotine exposition day onward. *** p < 0.001, ** p < 0.01, * p < 0.05.

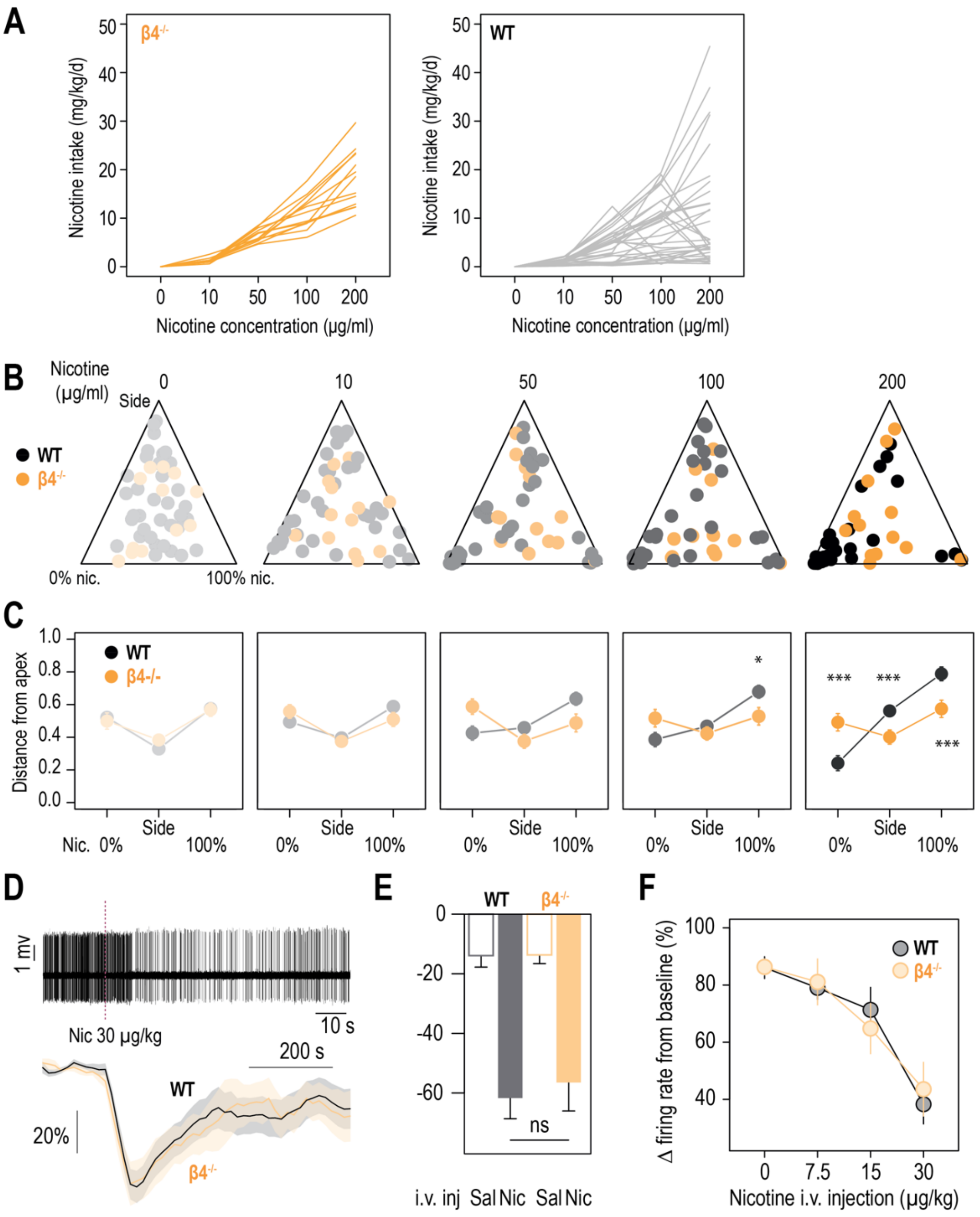


#### FIGURE SUPP 4

**Figure S4: Behavioral and electrophysiological differences between WT and β4$^{-/-}$ mice. A.** Nicotine intake (mg/kg/d) for individual WT (black) and β4$^{-/-}$ mice (orange), as concentration of nicotine increases. **B.** Ternary representations for WT (black) and β4$^{-/-}$ mice (orange), illustrating their nicotine consumption index over their side bias index, as nicotine concentration increases during the task. **C.** Distance from each of the three apices for WT (black) and β4$^{-/-}$ mice (orange) as nicotine concentration increases during the task. Note how the behavior of WT mice is highly nicotine-concentration dependent compared to that of β4$^{-/-}$ mice. (Mann-Whitney test, Nic 100 µg/ml : p(Nic) = 0.018; Nic 200 µg/ml : p(Sacc) < 0.001, p(Side) < 0.001, p(Nic) < 0.001) **D.** *In vivo* juxtacellular recordings of nicotine-evoked responses in nicotine-inhibited IPN neurons of WT and β4$^{-/-}$ animals. Top, representative electrophysiological recording of an IPN neuron, during an i.v. injection of nicotine (30 µg/kg). Bottom, average time course and amplitude of the change in firing frequency from baseline after an i.v. injection of saline and nicotine (30 µg/kg), for WT (black) and β4$^{-/-}$ animals (yellow). **E.** Average amplitude of the change in firing frequency from baseline after an i.v. injection of saline and nicotine (30 µg/kg) in nicotine-inhibited IPN neurons from WT (n =16 neurons from 14 mice) and β4$^{-/-}$ (n = 8 neurons from 6 mice) animals. **F.** Dose-dependent change in firing rate from baseline following i.v. injections of nicotine at 7.5, 15 and 30 mg/kg in nicotine-inhibited IPN neurons. *** p < 0.001, ** p < 0.01, * p < 0.05.

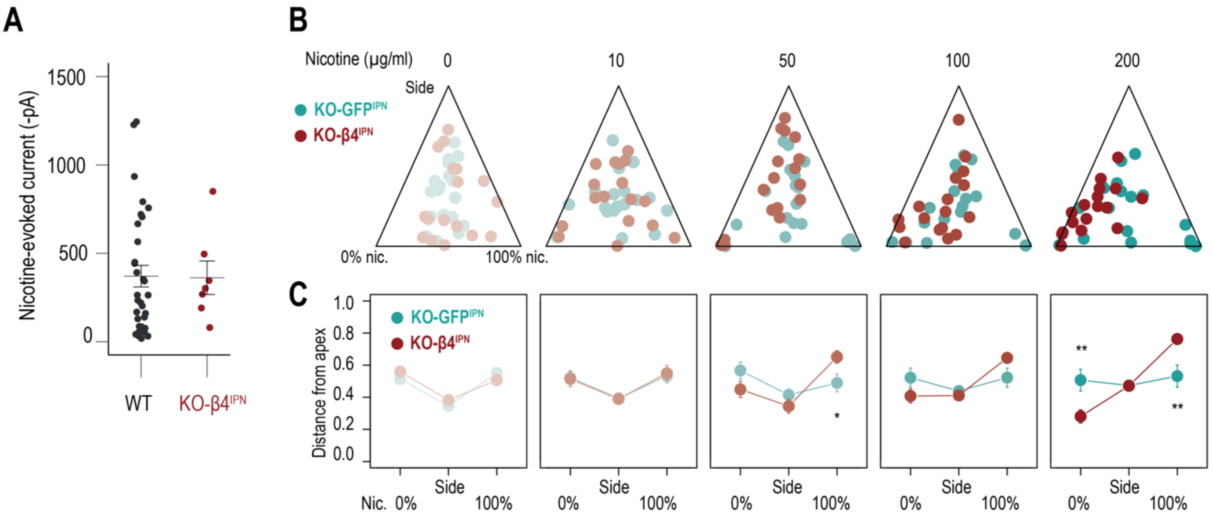


**FIGURE SUPP 5**

**Figure S5: Viral rescue of nAChR β4 subunit expression in the IPN of β4$^{-/-}$ mice.**

**A.** Average nicotine-induced currents following a puff application of nicotine (30 µM, 200 ms) on IPN neurons from WT (n = 32, -370 +/ -61 pA) and KO-β4$^{IPN}$ mice (n = 7, -362 ± 95 pA; Mann-Whitney, p = 0.6). **B.** Ternary representations for KO-GFP$^{IPN}$ (controls, green) and KO-β4$^{IPN}$ mice (red), illustrating their nicotine consumption index over their side bias index, as nicotine concentration increases during the task. **C.** Distance from each of the three apices for KO-GFP$^{IPN}$ (controls, green) and KO-β4$^{IPN}$ mice (red) as nicotine concentration increases during the task. (Mann-Whitney, Nic 50 µg/ml: p(Nic) = 0.017; Nic 200 µg/ml: p(Sacc) = 0.0074, p(Nic) = 0.0083). *** $p < 0.001$, ** $p < 0.01$, * $p < 0.05$.
